# Induction of hyperandrogenism and insulin resistance differentially modulates ferroptosis in uterine and placental tissues of pregnant rats

**DOI:** 10.1101/2020.03.30.015529

**Authors:** Yuehui Zhang, Min Hu, Wenyan Jia, Guoqi Liu, Jiao Zhang, Bing Wang, Juan Li, Peng Cui, Xin Li, Susanne Lager, Amanda Nancy Sferruzzi-Perri, Yanhua Han, Songjiang Liu, Xiaoke Wu, Mats Brännström, Linus R Shao, Håkan Billig

**Author notes:** Contributed equally to this work (Yuehui Zhang, Min Hu). Corresponding author: Linus R Shao, M.D., Ph.D.

## Abstract

Ferroptosis, a form of regulated necrotic cell death, plays roles in diverse physiological processes and diseases. Women with polycystic ovary syndrome (PCOS) have hyperandrogenism and insulin resistance (HAIR) and an increased risk of miscarriage and placental dysfunction during pregnancy. However, whether maternal HAIR alters mechanisms leading to ferroptosis in the gravid uterus and placenta remains unknown. Previous studies in rats showed that maternal exposure to 5α-dihydrotestosterone (DHT) and insulin (INS) from gestational day 7.5 to 13.5 induces HAIR and subsequently leads to placental insufficiency and fetal loss. We therefore hypothesized that maternal HAIR triggers ferroptosis in the uterus and placenta in association with fetal loss in pregnant rats. Compared with controls, we found that co-exposure to DHT and INS led to decreased levels of Gpx4 and glutathione (GSH), increased GSH+glutathione disulfide (GSSG) and malondialdehyde (MDA), aberrant expression of ferroptosis-associated genes (*Acsl4, Tfrc, Slc7a11*, and *Gclc*), increased iron deposition, and activated ERK/p38/JNK phosphorylation in the gravid uterus. However, in the placenta, DHT and INS exposure only partially altered the expression of ferroptosis-related markers (e.g., region-dependent Gpx4, GSH+GSSG, MDA, *Gls2* and *Slc7a11* mRNAs, and phosphorylated p38 levels). In the uteri co-exposed to DHT and INS, we also observed shrunken mitochondria with electron-dense cristae, which are key features of ferroptosis-related mitochondrial morphology, as well as increased expression of *Dpp4*, a mitochondria-encoded gene responsible for ferroptosis induction. In contrast, in placentas co-exposed to DHT and INS we found decreased expression of *Dpp4* mRNA and increased expression of *Cisd1* mRNA (a mitochondria-encoded iron-export factor). Further, DHT+INS-exposed pregnant rats exhibited decreased apoptosis in the uterus and increased necroptosis in the placenta. Our findings suggest that maternal HAIR causes the activation of ferroptosis in the gravid uterus and placenta, although this is mediated via different mechanisms operating at the molecular and cellular levels. Furthermore, our data suggest other cell death pathways may play a role in coordinating or compensating for HAIR-induced ferroptosis when the gravid uterus and placenta are dysfunctional.

**Suppl Figures:** https://doi.org/10.6084/m9.figshare.11794059.v5

## Introduction

Polycystic ovary syndrome (PCOS) is a complex and heterogeneous hormone-imbalance gynecological disorder that is influenced by genetic, environmental, and metabolic factors (1). This disorder affects approximately 4%–21% of all adolescent and reproductive-aged women and has a significant impact on their reproduction (2). Women with PCOS often suffer from hyperandrogenism (androgen excess) and insulin resistance (HAIR), and they are at high risk for miscarriage and obstetric complications during pregnancy (3, 4). Therapeutic interventions for different phenotypes and disease-related pregnancy complications in women with PCOS present a significant unmet medical need (5). Although it is thought that maternal, placental, and fetal defects all contribute to the onset and progression of miscarriage in PCOS patients, the pathogenesis of the pregnancy loss induced by HAIR and its precise regulatory mechanisms are still significant issues to be solved.

Ferroptosis is a recently described, iron-dependent form of regulated necrosis induced by oxidative stress, and it is distinct from other established forms of cell death such as apoptosis and necroptosis due to its unique morphological and biochemical features (6, 7). Growing evidence indicates that excessive or impaired ferroptosis plays a causative role in a variety of pathological conditions and diseases (8-11). It appears that the outcome of ferroptosis is programmed cell death, but which specific physiological processes or pathological conditions and disorders lead to ferroptosis activation remain poorly explored. The major molecular mechanisms and signaling pathways that are involved in the regulation of ferroptosis have been demonstrated in *in vivo* and *in vitro* studies (12). For example, suppression of glutathione (GSH) biosynthesis and subsequent inhibition or degradation of glutathione peroxidase 4 (Gpx4) activity, disturbed balance of iron homeostasis, and activation of the mitogen-activated protein kinase (MAPK) signaling pathways all contribute to regulate the initiation and execution of ferroptosis (6, 7, 10, 11). However, little is known about the role of ferroptosis (13) in comparison with other forms of programmed cell death such as apoptosis (14, 15) in female reproduction.

Given the significant association of PCOS with miscarriage during pregnancy (16, 17), we have recently demonstrated that the effects of HAIR in causing fetal loss is the consequence of uterine and placental defects (18, 19). To this end, we exposed pregnant rats to 5α-dihydrotestosterone (DHT) and insulin (INS) from gestational day (GD) 7.5 to 13.5 and found that this triggered many features of pregnant PCOS patients (including HAIR), as well as increased fetal loss. Moreover, the fetal loss was related to disrupted reactive oxygen species (ROS) production in the uterus and placenta of rat dams with induced HAIR. In particular, our previous animal experiments have shown that maternal HAIR-induced fetal loss is also associated with the inactivation of antioxidative proteins in the gravid uterus and placenta, namely nuclear factor erythroid 2-related factor 2 (Nrf2) and superoxide dismutase 1 (18, 19), which play an inhibitory role in the ferroptosis pathway (6, 10). Moreover, the expression of several other negative regulators of ferroptosis such as *Ho1* (heme oxygenase 1) (6) and *Mt1g* (metallothionein 1G) (20) are downregulated in the gravid uterus after combined maternal exposure to DHT and INS (18). Increased circulating ROS levels have been observed in both non-pregnant and pregnant rodents in which PCOS features have been induced (19, 21). Elevated ROS production and decreased anti-oxidative capacity has been observed in the ovarian granulosa cells and leukocytes of PCOS patients (21-23), and oxidative stress is proposed to contribute to miscarriage and infertility in women with PCOS (24, 25). It is therefore likely that the promotion of pathologic oxidative stress and activation of ferroptosis in the gravid uterus and placenta contribute to HAIR-induced fetal loss in both animal models and humans.

Mitochondria play a protective role in the regulation of exhausted GSH-induced ferroptosis (26, 27). In women with PCOS and miscarriage (24, 28), as well as in pregnant PCOS-like rodents with fetal loss (18, 19, 29, 30), there is mounting evidence for mitochondrial abnormalities and oxidative damage. For instance, decreased mitochondrial DNA copy number is associated with the developmental clinical phenotype and severity of PCOS, and several mitochondria-tRNA mutations are seen in PCOS patients. In addition, aberrant expression of mitochondrial biogenesis genes, oxidative phosphorylation and anti-oxidative proteins are found in PCOS patients who suffer from recurrent miscarriage, as well as in PCOS-like rodents. On the basis of these preclinical and clinical studies, we hypothesized that maternal HAIR triggers impairments in Gpx4/GSH-regulated lipid peroxidation and iron-associated and mitochondria-mediated ferroptosis in the gravid uterus and placenta resulting in increased fetal loss during pregnancy.

The aim of this study was to determine whether exposure to DHT and INS in pregnant rats (which induces HAIR/PCOS (18, 19)) leads to activation of the ferroptosis cascade and malondialdehyde (MDA, a marker of oxidative stress), iron accumulation, and perturbed mitochondrial function in the uterus and placenta. Further, we conducted a parallel analysis of the expression of genes and proteins that are involved in necroptosis and apoptosis, two other programmed cell death pathways that might contribute to defects in the gravid uterus and the placenta. This study is the first to report an association between HAIR and different forms of regulated cell death in the gravid uterus and placenta *in vivo*. Our findings indicate that ferroptosis is one of the potential mechanisms by which maternal HAIR leads to uterine and placental dysfunction and at least partially explains the resultant fetal loss observed.

## Materials and Methods

### Ethics approval

All experiments were conducted in compliance with all relevant local ethical regulations. Animal experiments were approved and authorized by the Animal Care and Use Committee of the Heilongjiang University of Chinese Medicine, China (HUCM 2015-0112), and followed the National Institutes of Health guidelines on the care and use of laboratory animals. The human study protocol conformed to the principles outlined in the Declaration of Helsinki under approval from the institutional Ethical Review Committee of the Obstetrics and Gynecology Hospital of Fudan University, Shanghai, China (OGHFU 2013-23), and informed consent was obtained from each patient in written form.

### Animals, experimental setting, and tissue collection

Adult Sprague–Dawley female and male rats were obtained from the Laboratory Animal Centre of Harbin Medical University, Harbin, China. All animals were health checked daily throughout the experiment and were maintained in an environmentally controlled and pathogen-free barrier facility on a standard 12 h light/dark cycle at 22 ± 2°C and 55–65% humidity and with free access to normal diet and water. Before the experiment, female rats were allowed to acclimatize for a minimum of 7 days and then were monitored daily by vaginal lavage to determine the stage of the estrous cycle (31, 32). Pregnancy was achieved by housing female rats on the night of proestrus with fertile males of the same strain at a 2:1 ratio. Confirmation of mating was defined by the presence of a vaginal plug, and this was considered as GD 0.5. The rats were sacrificed between 0800 and 0900 hours on GD 14.5. All animal procedures in this study were performed as described in our previous publications (18, 19).

To induce HAIR, pregnant rats were randomly assigned to be intraperitoneally injected with DHT (1.66 mg/kg/day, suspended in sesame oil, Sigma-Aldrich, St. Louis, MO, USA) and/or human recombinant INS (6.0 IU/day, diluted in sterile saline, Eli Lilly Pharmaceuticals, Giza, Egypt) or an equal volume of saline and sesame oil as controls on GD 7.5 as previously described (18, 19). This therefore generated the following four study groups: Control, DHT+INS, DHT, and INS. All animals were treated for 7 consecutive days. The dose of DHT used in our rats was chosen to mimic the hyperandrogenic state in PCOS patients who have approximately 1.7-fold higher circulating DHT concentrations compared to healthy controls (33, 34). The dose of INS was chosen because it induces metabolic disturbances including peripheral and uterine insulin resistance in rats (32, 35). The body weight, oral glucose tolerance test and circulating levels of androgens (testosterone, dehydroepiandrosterone, and DHT) were measured to confirm that HAIR was induced by exposure to DHT and INS, as reported previously (18, 19). Pregnant rats co-exposed to DHT and INS had metabolic and endocrine aberrations at GD 14.5, thus replicating the changes observed in pregnant PCOS patients (36, 37). On GD 14.5, tissues, including the maternal uterus and placenta, as well as fetuses were dissected. These were then either fixed for morphological and immunohistochemical analyses or immediately frozen in liquid nitrogen and stored at −70°C for quantitative real-time polymerase chain reaction (qPCR) (38) and Western blot analyses. Only viable conceptuses (fetuses and placentas) were analyzed further.

### Gene expression analysis by qPCR

The isolation and quantification of the RNA and the qPCR assay were performed as previously described (32, 39). The PCR amplifications were performed with SYBR green qPCR master mix (#K0252, Thermo Scientific, Rockford, IL). Total RNA was prepared from the frozen whole uterine and placental tissues, and single-stranded cDNA was synthesized from each sample (2 μg) with M-MLV reverse transcriptase (#0000113467, Promega Corporation, Fitchburg, WI) and RNase inhibitor (40 U) (#00314959, Thermo Scientific). Uterine and placental RNA purities (A260/A280 ratios) were evaluated with a NanoDrop 1000 spectrometer (Thermo Fisher Scientific). Only samples presenting a ratio greater than 1.8 were kept for further analyses. The integrity of the extracted RNA samples was additionally determined by using an Experion RNA StdSens Analysis Kit (Bio-Rad). Any samples showing poor RNA quality were also excluded from further analysis. cDNA (1 μl) was added to a reaction master mix (10 μl) containing 2× SYBR green qPCR reaction mix (Thermo Scientific) and gene-specific primers (5 μM of forward and reverse primers). All reactions were performed at least twice, and each reaction included a non-template control, and specific sample sizes are denoted in the figure legends. Fold changes in mRNA expression were calculated by the ΔΔCT method using *Gapdh* (18) as the reference gene, which was stably expressed between the groups (38). Results are expressed as fold changes after normalizing to the control group. The qPCR primers used in this study are listed in **Table 1**. All sets of primers were validated for qPCR prior to analysis. This involved determining that the efficiency of amplification using a standard curve of cDNA was above 85% and not different from the *Gapdh* reference gene, and there were no non-specific PCR products seen in a melt curve analysis immediately after the amplification or in parallel reactions with un-transcribed RNA or in reactions without templates (the negative controls). Further, in order to avoid introducing variability, all uterine and placental samples for a given target gene were analysed on a single plate.

**Table 1.**
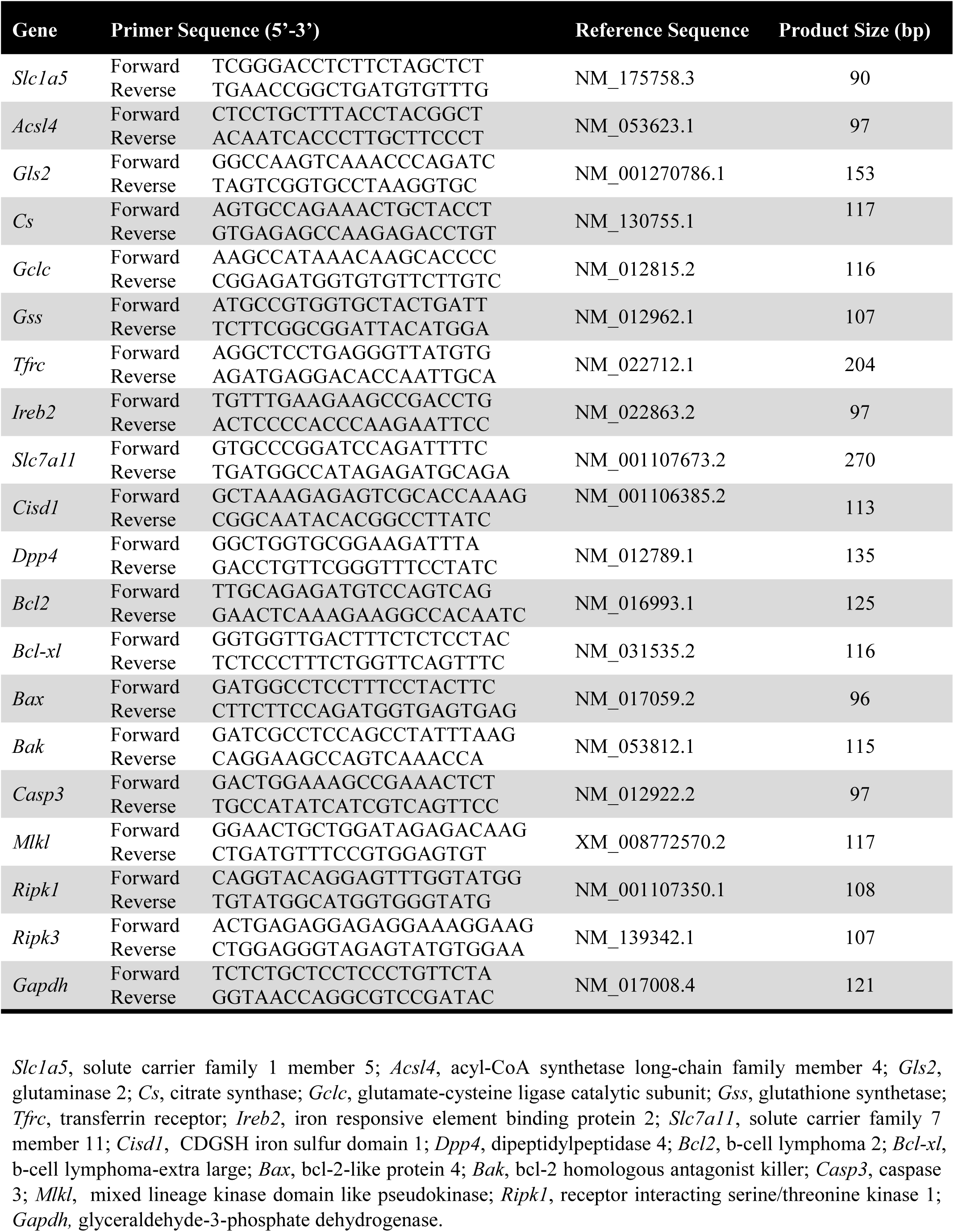
Primer sequences used for qPCR measurement.

### Protein isolation and Western blot analysis

A detailed explanation of the tissue lysate preparation and the Western blot analysis protocol has been published (32, 39). Tissue proteins were isolated by homogenization in RIPA buffer (Sigma-Aldrich) supplemented with cOmplete Mini protease inhibitor cocktail tablets (Roche Diagnostics, Mannheim, Germany) and PhosSTOP phosphatase inhibitor cocktail tablets (Roche Diagnostics). After determining the total protein concentration by Bradford protein assay (Thermo Fisher Scientific), equal amounts (30 μg) of protein were resolved on 4–20% TGX stain-free gels (Bio-Rad Laboratories GmbH, Munich, Germany) and transferred onto PVDF membranes. The membranes were probed with anti-Gpx4 antibody (ab125066, Abcam, Cambridge, UK), anti-ERK1/2 antibody (#4695, Cell Signaling Technology, Danver, MA, USA), anti-phospho-ERK1/2 antibody (#9911, Cell Signaling Technology), anti-p38 MAPK antibody (#8690, Cell Signaling Technology), anti-phospho-p38 MAPK antibody (#4511, Cell Signaling Technology), anti-JNK antibody (#9252, Cell Signaling Technology), anti-phospho-JNK antibody (#4668, Cell Signaling Technology), and anti-cleaved caspase-3 antibody (#9664, Cell Signaling Technology) all diluted 1:1,000 in 0.01 M Tris-buffered saline supplemented with Triton X-100 (TBST) containing 5% w/v non-fat dry milk followed by anti-rabbit IgG HRP-conjugated goat secondary antibody (A0545, Sigma-Aldrich). Signal was detected using the SuperSignal West Dura Extended Duration Substrate (Thermo Fisher Scientific) and captured using a ChemiDoc MP Imaging System (Bio-Rad). Initial experiments were performed to verify the identification of cytosolic, mitochondrial and nuclear Gpx4 by Western blot analysis using rat testis (which has high expression compared to other tissues (40, 41)), epididymis and ovary (**Supplemental Fig. 1)**. For each Western blot, ultraviolet activation of the Criterion stain-free gel was used to assess total protein loading for each sample (32). Band densitometry was performed using Image Laboratory (Version 5.0, Bio-Rad) and the intensity of each protein band was normalized to the total protein in the individual sample. Due to the number of samples per group, multiple gels were run per group, each containing three replicates per group. For quantification and to ensure standardisation across blots, the expression of the target protein was normalised to the mean value for the control group on the blot and then all the normalised values were statistically compared to assess the effect of the treatment groups. This ensured that we could accurately compare protein abundance across groups with the one tissue.

### Histological processing and Gpx4 immunohistochemistry

Histological processing and immunohistochemistry were performed according to previously described methods (32, 42). Fresh tissues were dissected and immediately fixed in 4% formaldehyde in neutral buffered solution at 4°C for 24 h and then embedded in paraffin. Sections (5 μm) were deparaffinized and rehydrated in xylene and graded series of ethanol (99.99%, 80%, and 70% in distilled water, Sigma-Aldrich) for 10 min each. After incubation with the Gpx4 antibody (1:200 dilution, Abcam) overnight at 4°C in a humidified chamber, the sections were stained using the avidin-biotinylated-peroxidase ABC kit followed by a 5-min treatment with 3,3’-diaminobenzidine (DAB, SK-4100, Vector Laboratories). Stained sections were observed and imaged on a Nikon E-1000 microscope (Japan) under bright-field optics and photomicrographed using Easy Image 1 (Bergström Instrument AB, Sweden). A negative control was performed by using the same concentration of isotype-matched rabbit IgG instead of the primary antibody. Only minimal cytoplasmic background staining was observed (**Supplemental Fig. 2 *right***).

### Perls’ histochemical reaction

Iron deposition was detected using DAB-enhanced Perls’ staining as previously described (43). After deparaffinization and rehydration, sections were immersed in a mixture of equal volumes of potassium ferrocyanide solution (HT201, Sigma-Aldrich) and hydrochloric acid solution (HT201, Sigma-Aldrich) for 1 h at room temperature. Sections were washed with PBS five times for 5 min each and incubated with DAB for 10 min and pararosaniline solution (HT203, Sigma-Aldrich) for 2 min. Images of excess iron deposits were captured on a Nikon E-1000 microscope (Japan) under bright-field optics and photomicrographed using Easy Image 1 (Bergström Instrument AB, Sweden).

### Mitochondria structure by transmission electron microscopy (TEM)

TEM was performed according to a published method (30). Fresh uterine and placental tissues were fixed in 2.5% glutaraldehyde in phosphate buffered saline (PBS, pH 7.2–7.4) for 1 h at room temperature (RT) and further rinsed with 0.1 M PBS three times for 15 minutes each. Secondary fixation with 1% osmium tetroxide in PBS was performed for 1 h prior to sequential dehydration with an acetone gradient (50%, 70%, and 90% for 15 min each and 100% three times for 30 min each time at RT). Samples were finally embedded in Epon epoxy resin. Random areas from uterine and placental tissues were oriented for ultrastructural analysis. The blocks were cut in 50–60 nm sections using a Reichert ultramicrotome (Leica, Germany), collected on 300 mesh copper grids, and stained with 3% uranyl acetate and counterstained with lead citrate before visualization. The post-stained sections were examined and imaged with a transmission electron microscope (H-7650, Hitachi, Japan) equipped with an electron imaging spectrometer. Image collection and parameter settings were identical for each of the different tissues/regions analyzed.

### Quantification of GSH, MDA, and mitochondrial open reading frame of the 12S rRNA-c (MOTS-c)

The intracellular GSH, MDA, and MOTS-c levels were assessed using a GSH/GSH+glutathione disulfide (GSSG) assay kit (ab239709, Abcam, Cambridge, UK), MDA assay kit (ab118970, Abcam), and MOTS-c ELISA kit (CEX132Ra, Cloud-Clone/USCNK, Oxfordshire, UK), respectively, according to the manufacturers’ protocols. A standard curve for GSH, MDA, and MOTS-c concentration was generated and used for calculating their concentration in the samples. The concentration of GSH, GSH+GSSG, MDA, and MOTS-c in each group was normalized to the total tissue protein concentration as determined by the Bradford protein assay (Thermo Fisher Scientific).

### Data processing, statistical analysis, and graphical representations

No statistical methods were used to predetermine the sample size. Data are presented as the means ± SEM, and the sample size (n) is listed in the figure legends and indicates the number of animals in each experiment. Statistical analyses were performed using SPSS version 24.0 for Windows (SPSS Inc., Chicago, IL). The normal distribution of the data was tested with the Shapiro– Wilk test. Differences between groups were analyzed by one-way ANOVA followed by Tukey’s post-hoc test for normally distributed data or the Kruskal–Wallis test for skewed data (**Supplemental Table 1**). All p-values less than 0.05 were considered statistically significant.

## Results

Because we were most interested in how HAIR induces changes in ferroptosis as opposed to apoptosis and necroptosis in gravid uterine and placental tissues, we mainly describe the observations in DHT+INS-exposed pregnant rats vs. control pregnant rats.

### Differential regulation of Gpx4 and GSH protein expression in the gravid uterus and placenta exposed to DHT and INS

Gpx4 is present in the cytoplasm, mitochondria and nucleus in mammalian cells (44) and in our Western blot analysis, the ∼20-kDa band represents the cytosolic and mitochondrial Gpx4 protein, whereas the ∼34-kDa band represents the nuclear Gpx4 protein (**Supplemental Fig. 1**). We initially performed Western blot and immunohistochemical analyses to characterize the tissue and intracellular localization of Gpx4 protein in rat and human uterine and placental tissues. The Western blot analysis revealed a predominant band corresponding to cytosolic and mitochondrial Gpx4 (under denaturing and reducing conditions) in the diestrus uterus of rats and in the non-pregnant secretory phase and early pregnant decidualized endometria of humans (**Supplemental Fig. 3A**). Our further immunohistochemical studies showed that while positive immunostaining for cytosolic Gpx4 was mainly observed in luminal and glandular epithelial cells, Gpx4 immunostaining was additionally localized to the nucleus of non-decidualized and decidualized stromal cells and myometrial smooth muscle cells in rats and humans (**Supplemental Fig. 3B-D**). This is consistent with variations in the cellular compartmentalization of Gpx4 in the cytosol and nucleus between different cell types (44). In control pregnant rats, immunohistochemical analysis showed that Gpx4 was localized to both the cytosol and nucleus of different cell types in the gravid uterine decidua, myometrium and placenta (**Fig. 1C1-C4** and **Supplemental Fig. 2B**). Although the significance of mitochondrial and nuclear Gpx4 remains to be determined (45), cytosolic Gpx4 has been identified as a central regulator of ferroptosis (11). We thus evaluated Gpx4 expression and localization in the gravid uterus and placenta exposed to DHT and INS. The Western blot analysis revealed a significant decrease in uterine Gpx4 protein abundance in DHT+INS-exposed pregnant rats (**Fig. 1A**). Consistent with this, we found weaker immunoreactivity of Gpx4 in the cytosolic compartments of decidualized stromal cells and smooth muscle cells in the gravid uterus of DHT+INS-exposed pregnant rats (**Fig. 1D1-D2)**. Although there was no significant reduction in uterine Gpx4 protein abundance by Western blot analysis in the pregnant rats treated alone with DHT or INS (**Fig. 1A**), fewer cells of the gravid uterus showed nuclear immunostaining of Gpx4 when compared to controls (**Fig. 1B**). Although Gpx4 protein abundance by Western blot analysis was not significantly reduced in the placenta of DHT+INS-exposed pregnant rats (**Fig. 1A**), immunohistochemical analysis revealed that the level of Gpx4 immunostaining was lower in trophoblast populations of the placenta in DHT+INS-exposed pregnant rats compared to the control pregnant rats (**Fig. 1D3-D4**). In particular, Gpx4 immunostaining was no longer localized to the nuclei of spongiotrophoblasts and glycogen cells of DHT+INS-exposed pregnant rats (**Fig. 1E3-E4**). There was no alteration of placental Gpx4 protein abundance in pregnant rats treated with DHT alone or INS alone (**Fig. 1A**). However, similar to what was seen in DHT+INS-exposed pregnant rats, Gpx4 was no longer localized to the nuclei of spongiotrophoblasts and glycogen cells in pregnant rats treated with DHT alone (**Fig. 1E3**) but not INS alone (**Fig. 1F3**).

**Figure 1.**
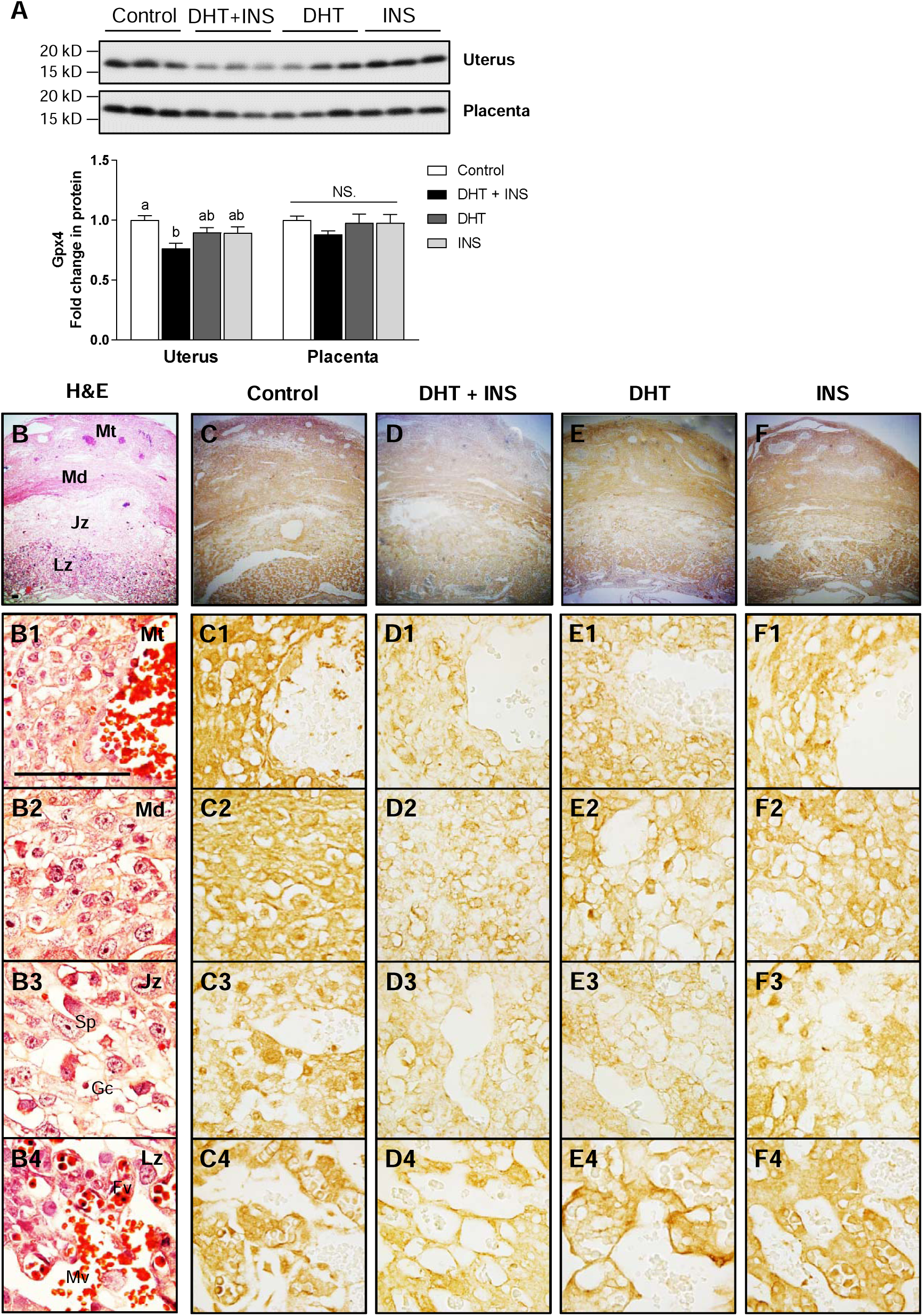
Regulation and localization of Gpx4 protein in pregnant rats exposed to DHT and/or insulin at GD 14.5. Western blot analysis of Gpx4 protein expression in the uterus and placenta (A, n = 9/group). In all plots, values are expressed as means ± SEM. Significant differences (p < 0.05) within each group are denoted by different letters, and the same letter between groups indicates lack of statistical significance. N.S., not significant. Tissue sections were stained with hematoxylin and eosin (H&E, B1-4). Histological analysis by Gpx4 immunostaining in the gravid uterus (Mt and Md) and placenta (Jz and Lz) (C1-F4). Images are representative of 8−10 tissue replicates per group. Mt, mesometrial triangle; Md, mesometrial decidua; Jz, junctional zone (maternal side); Gc, glycogen cells; Sp, spongiotrophoblast cells; Lz, labyrinth zone (fetal side); Mv, maternal blood vessel; Fv, fetal blood vessel. Scale bars (100 μm) are indicated in the photomicrographs. DHT, 5α-dihydrotestosterone; INS, insulin.

Because Gpx4 uses GSH as a substrate in its peroxidase reaction cycle (46) and because GSH depletion is one of the key triggers for ferroptosis (10, 11), we measured the levels of GSH and GSH+GSSG in the gravid uterus and placenta exposed to DHT and INS. As shown in **Figure 2A**, co-exposure to DHT and INS resulted in decreased GSH levels in the gravid uterus, but not in the placenta, while increased GSH+GSSG levels were detected in both tissues. Similarly, alterations of GSH and GSH+GSSG levels were found in the gravid uterus exposed to insulin alone. Levels of GSH were lower in both the gravid uterus and placenta exposed to DHT alone compared to controls (**Fig. 2A**).

**Figure 2.**
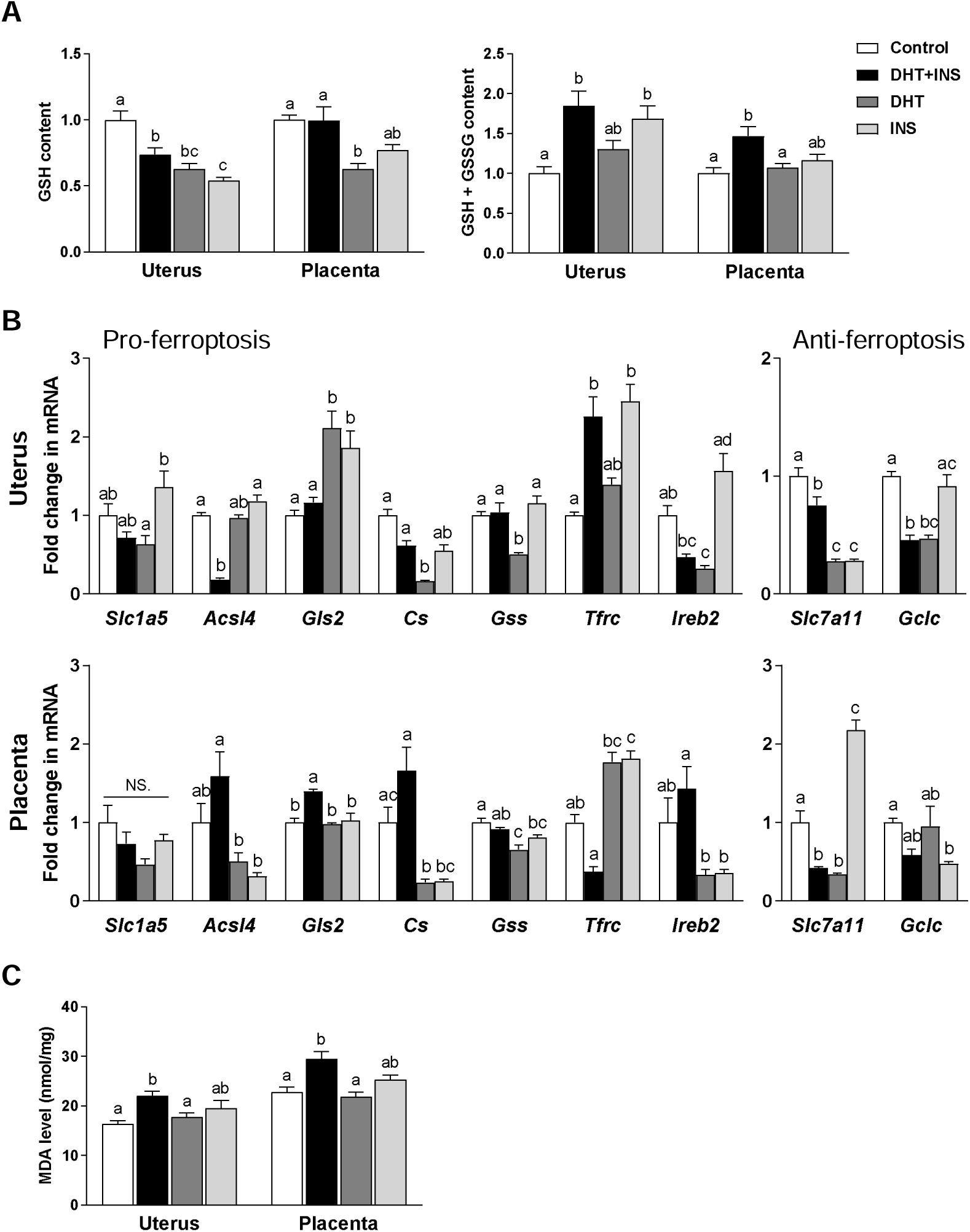
Alteration of GSH, GSH+GSSG, ferroptosis-related gene expression, and MDA in pregnant rats exposed to DHT and/or insulin at GD 14.5. ELISA analysis of GSH, GSH+GSSG, and MDA in the uterus and placenta (A, n = 8/group). qPCR analysis of uterine and placental genes involved in modulating ferroptosis (B, n = 7−8/group). In all plots, values are expressed as means ± SEM. Significant differences (p < 0.05) within each group are denoted by different letters, and the same letter between groups indicates lack of statistical significance. DHT, 5α-dihydrotestosterone; INS, insulin.

### Alterations of ferroptosis-related gene expression, MDA levels, intracellular iron deposition, and the MAPK signaling pathway

Next, we examined whether maternal exposure to DHT and/or INS alters the expression of pro-ferroptosis (*Slc1a5, Acsl4, Gls2, Cs, Gss, Tfrc* and *Ireb2*) or anti-ferroptosis (*Slc7a11*, and *Gclc*) genes in the gravid uterus and placenta. In pregnant DHT+INS-exposed rats, uterine *Acsl4, Slc7a11* and *Gclc* mRNAs were decreased, while *Tfrx* mRNA was increased (**Fig. 2B**). In comparison with the control uterus, maternal exposure to DHT alone decreased *Cs, Gss, Ireb2, Slc7a11* and *Gclc* mRNA expression, whereas exposure to INS alone increased *Gls2* and *Tfrc* mRNAs and decreased *Slc7a11* mRNA expression (**Fig. 2B**). qPCR analysis also showed that *Gls2* mRNA expression was increased and *Slc7a11* mRNA expression was decreased in the placenta after co-exposure to DHT and INS. In comparison with the control placenta, exposure to DHT alone decreased *Cs, Gss* and *Slc7a11* mRNA expression, whereas exposure to INS alone decreased *Gss* and *Gclc* mRNAs in parallel to increased *Tfrc* and *Slc7a11* mRNA expression (**Fig. 2B**).

Given that one of the key consequences of ferroptosis is elevated lipid peroxidation (6, 7), we next examined the impact of DHT and INS on the levels of MDA, a marker of lipid peroxidation (47), in the gravid uterus and placenta. As shown in **Figure 2C**, co-exposure to DHT and INS resulted in increased MDA levels in both the gravid uterus and placenta. However, there were no significant changes in MDA levels between the DHT-exposed rats or the INS-exposed rats and control rats.

Whether chronic exposure to DHT and INS can modulate tissue iron deposition was also examined. Perls’ histochemical reaction showed specific cytoplasmic and granular iron storage in rat uterine epithelial and decidualized stromal cells on GD 6, which is prior to the induction of HAIR (**Supplemental Fig. 4A1-A2**). These data are consistent with a previous report showing the cellular expression of ferritin heavy chain, a component of the multi-subunit iron-binding protein ferritin, in the uterus during early pregnancy (48). In the gravid uterus, iron accumulation was increased in the external muscle layer, the mesometrial triangle, as well as in the decidua of DHT+INS-exposed pregnant rats (**Fig. 3B1-B3** and **Supplemental Fig. 4B3-B4**) compared to control pregnant rats (**Fig. 3A1-A3** and **Supplemental Fig. 4B1-B2**). Similarly, a significant increase in iron storage in the mesometrial triangle was also observed in DHT-exposed pregnant rats (**Fig. 3C1-C2**). Further, granular and cytoplasmic iron-positive staining was barely detectable in the mesometrial decidua in the DHT-exposed rats and in the INS-exposed rats (**Fig. C2 and D3**). No iron-positive staining was found in the placental junctional zone in any of the experimental groups (**Fig. 3A4, B4, C4, and D4**), while intense iron-positive staining was consistently detected in immature erythrocytes within the placental labyrinth zone in all experimental groups (**Fig. 3A5, B5, C5 and D5**). These results indicate that the amount of deposited iron was elevated, especially in the gravid uterus, following exposure to DHT and/or INS.

**Figure 3.**
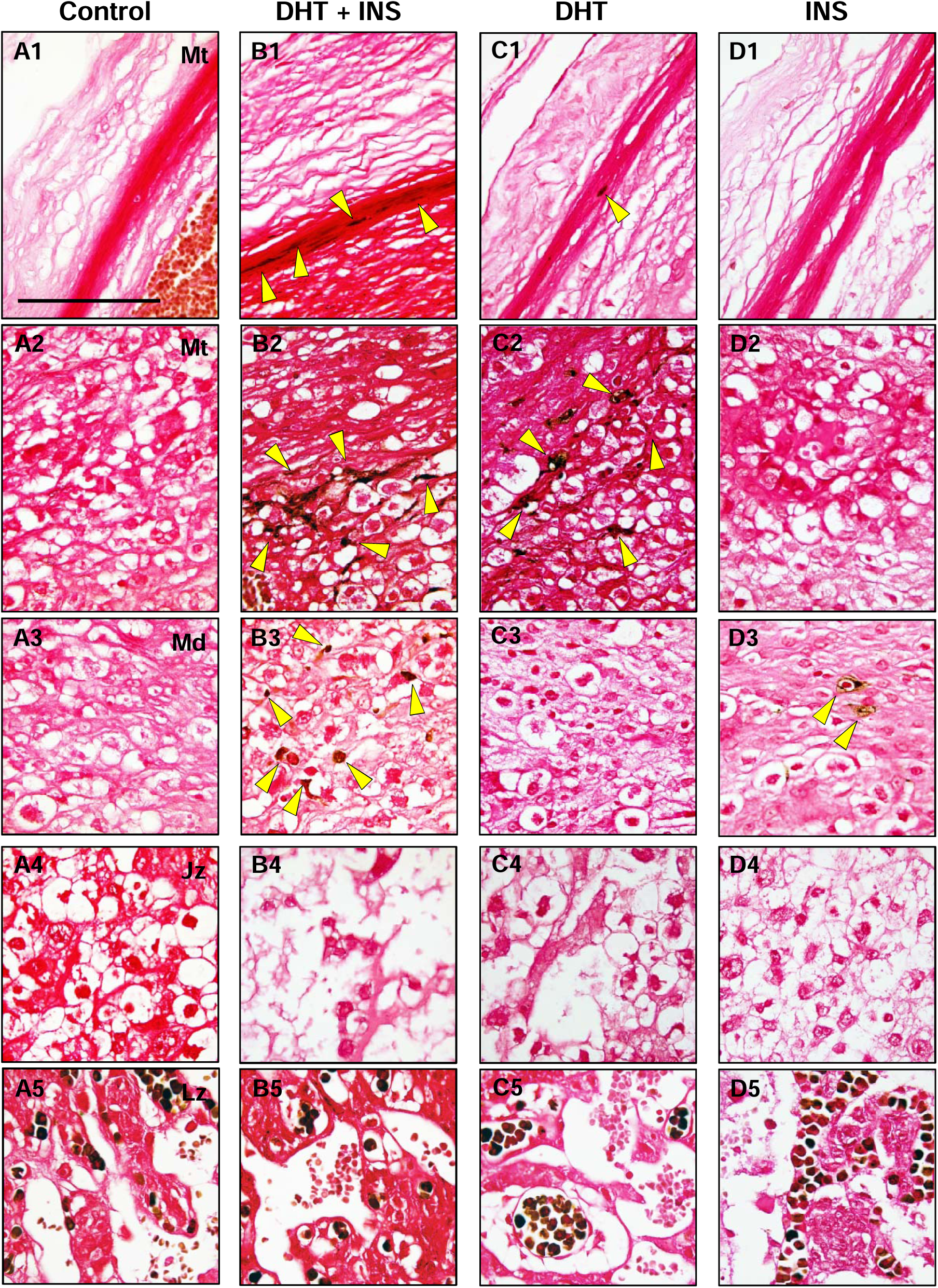
Iron deposition in the uterus and placenta of pregnant rats exposed to DHT and/or insulin at GD 14.5. Gravid uterine and placental tissues from pregnant rats treated with vehicle (A1-5), DHT+INS (B1-5), DHT (C1-5), or INS (D1-5) are shown. The sections were stained by DAB-enhanced Perls’ staining for iron accumulation. Yellow arrowheads indicate iron-positive staining. Images are representative of eight tissue replicates per group. Mt, mesometrial triangle; Md, mesometrial decidua; Jz, junctional zone (maternal side); Lz, labyrinth zone (fetal side). Scale bars (100 μm) are indicated in the photomicrographs. DHT, 5α-dihydrotestosterone; INS, insulin.

Taking into consideration that the MAPK signaling pathway, including ERK, p38, and c-JUN NH_2_-terminal kinase (JNK) is involved in the execution of ferroptosis in other cells (10, 49), we evaluated whether co-exposure to DHT and INS may be linked to activation of the MAPK signaling pathway in the gravid uterus and placenta. As shown in **Figure 4A**, in the gravid uterus DHT+INS exposure resulted in an increased abundance of phosphorylated ERK1/2 (p-ERK1/2) and decreased total ERK1/2, which subsequently resulted in an increased p-ERK1/2:ERK1/2 ratio. Moreover, both p-JNK and total JNK protein abundance were increased, whereas the p-JNK:JNK ratio remained unchanged in the DHT+INS-exposed gravid uterus compared to the control uterus (**Fig. 4A**). Additionally, a similar increase in p-p38 protein abundance and the p-p38:p38 ratio was observed in both the gravid uterus (**Fig. 4A**) and placenta (**Fig. 4B**) after co-exposure to DHT and INS. These results indicate that both ERK1/2 and JNK signaling are only activated in the gravid uterus, whereas p38 signaling is activated in both the gravid uterus and placenta after co-exposure to DHT and INS.

**Figure 4.**
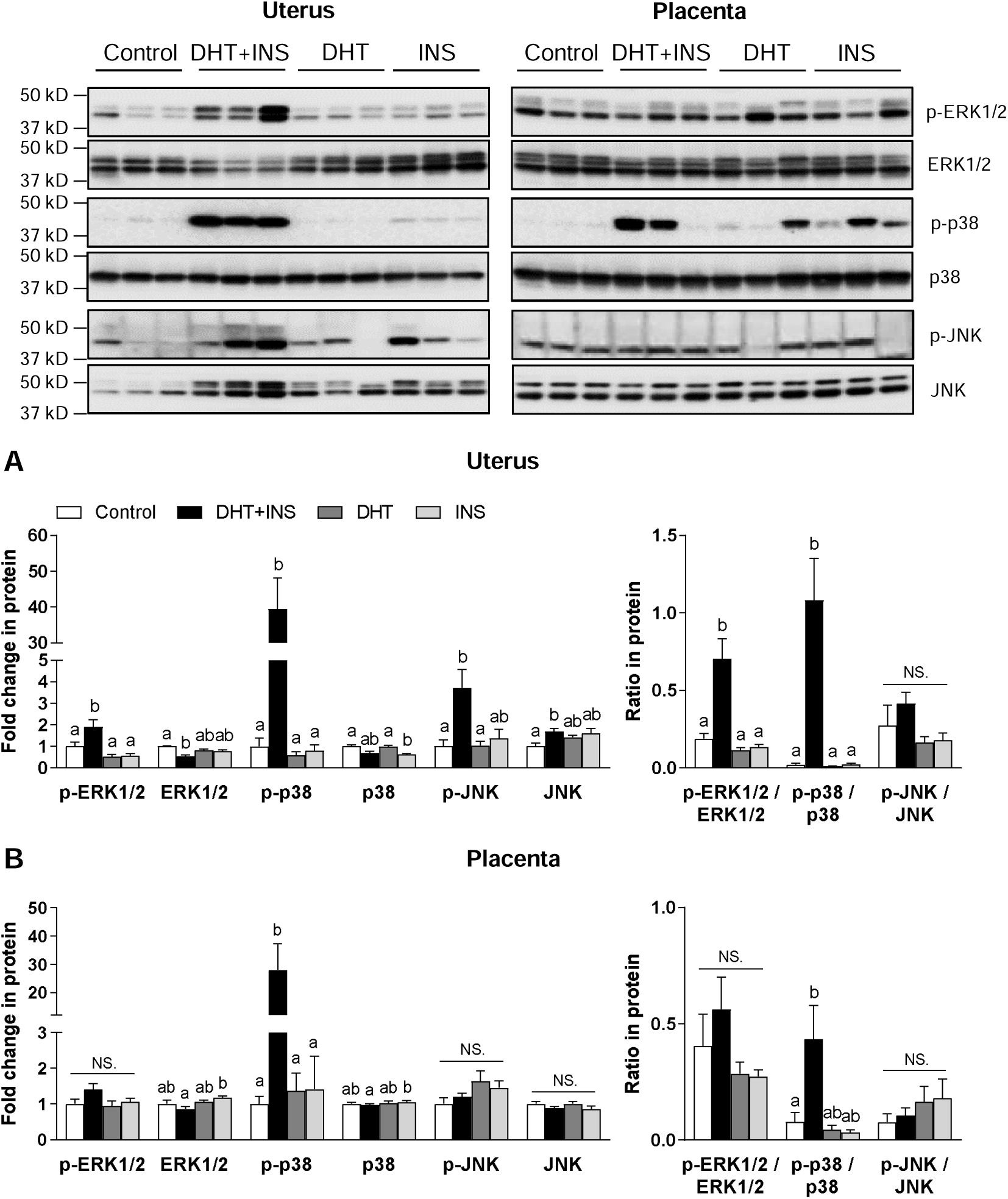
Changes in the expression of proteins involved in the ferroptosis-related MAPK signaling pathway in pregnant rats exposed to DHT and/or insulin at GD 14.5. Western blot analysis of ERK, p38, and JNK protein expression and their phosphorylated forms in the uterus and placenta (n = 9/group). In all plots, values are expressed as means ± SEM. Significant differences (p < 0.05) within each group are denoted by different letters, and the same letter between groups indicates lack of statistical significance. N.S., not significant. DHT, 5α-dihydrotestosterone; INS, insulin.

### Changes in mitochondrial morphology are associated with changes in mitochondria-encoded gene and protein expression

By TEM (**Supplemental Fig. 5**), we found that compared to controls (**Fig. 5A1**) the mitochondria showed swelling and collapsed and poorly defined tubular cristae in the gravid rat uterus exposed to DHT and/or INS (**Fig. 5B1-D1**). Consistent with ferroptosis-related mitochondrial morphology (7, 10, 12), shrunken mitochondria with numerous electron-dense cristae or with the absence of cristae were only found in the DHT+INS-exposed gravid uterus (**Fig. 5B1 *arrows***). TEM also revealed that treatment with DHT or INS reduced the number of mitochondrial cristae (**Fig. 5C1 and D1**). Further, ultrastructural analysis showed that mitochondria in the trophoblast within the junctional zone were significantly affected by exposure to DHT and/or INS (**Fig. 5A2-D2**). For instance, mitochondria showed blebbing, few or no tubular cristae, and decreased electron density in all treatment groups (**Fig. 5B2-D2**). However, there was less mitochondrial damage observed in the trophoblast of the labyrinth zone in all treatment groups compared to controls (**Fig. 5A3-D3**).

**Figure 5.**
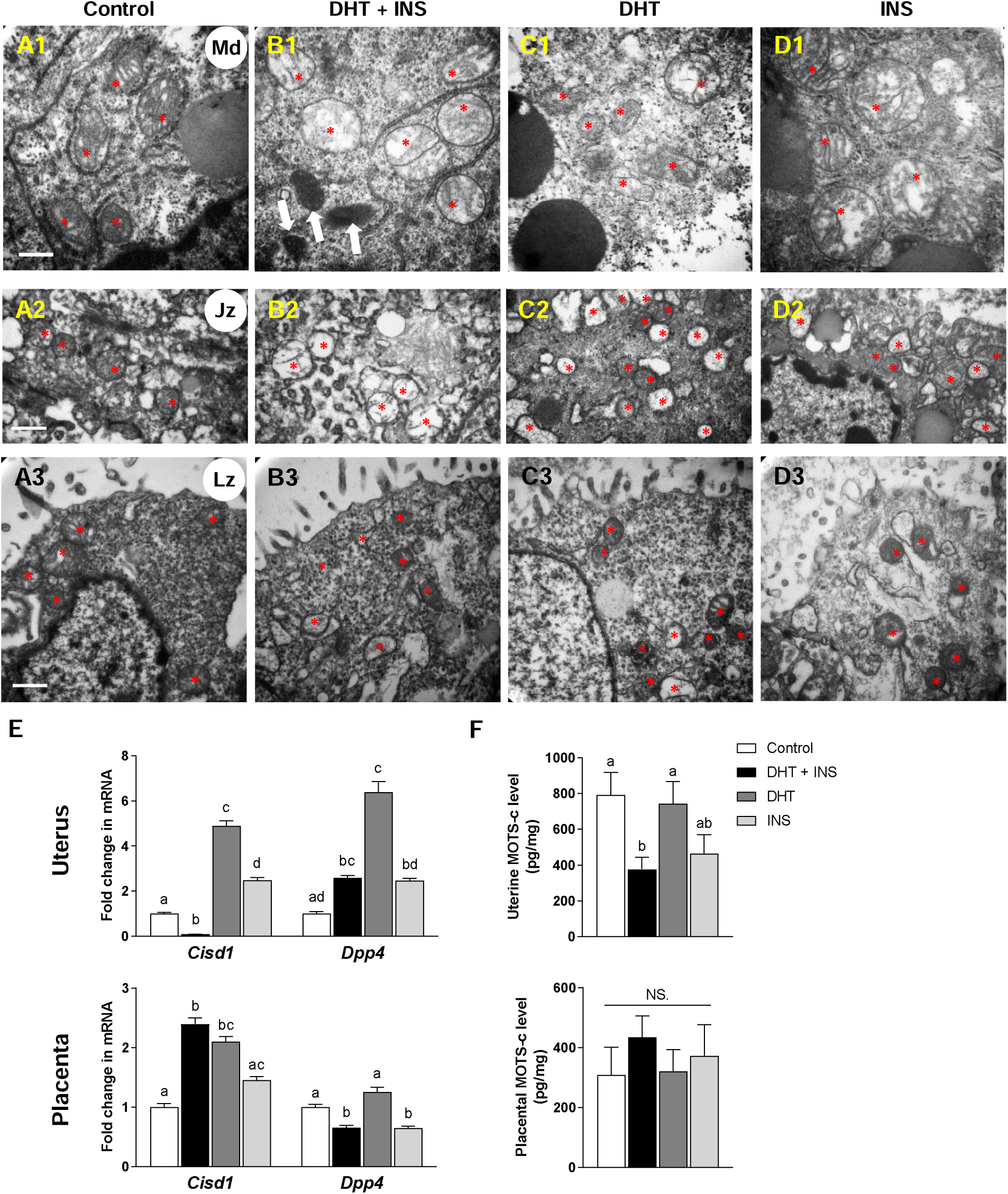
Electron microscopy and mitochondria-mediated ferroptosis-related gene and protein expression in pregnant rats exposed to DHT and/or insulin at GD 14.5. Mitochondrial ultrastructural defects in the uterus (A1, B1, C1, and D1, mesometrial decidua) and placenta (junctional (A2, B2, C2, and D2) and labyrinth zones (A3, B3, C3, and D3)). Images are representative of two tissue replicates. Md, mesometrial decidua; Jz, junctional zone (maternal side); Lz, labyrinth zone (fetal side). Red asterisks indicate mitochondria, and white arrows indicate shrunken mitochondria with electron-dense cristae. Scale bars (500 nm) are indicated in the photomicrographs. qPCR analysis of mitochondrial genes involved in modulating ferroptosis (E, n = 8/group). ELISA analysis of MOTS-c content (F, n = 8/group). In all plots, values are expressed as means ± SEM. Significant differences (p < 0.05) within each group are denoted by different letters, and the same letter between groups indicates lack of statistical significance. N.S., not significant. DHT, 5α-dihydrotestosterone; INS, insulin.

Based on these morphological observations, the expression of known mitochondria-encoded genes (*Cisd1*, an anti-ferroptosis gene, and *Dpp4*, a pro-ferroptosis gene (6, 11)) and protein (MOTS-c, an enhancer of insulin sensitivity (50)) was analyzed by qPCR (**Fig. 5E**) and ELISA (**Fig. 5F**). In the pregnant rat uterus, DHT+INS-exposure decreased *Cisd1* mRNA expression, increased *Dpp4* mRNA expression, and decreased the MOTS-c protein level (**Fig. 5E and F *upper panel***). In contrast, we found significantly higher uterine *Cisd1* and *Dpp4* mRNA expression in INS-exposed pregnant rats (**Fig. 5E *upper panel***), but unchanged uterine MOTS-c protein levels in DHT-exposed pregnant rats compared to controls (**Fig. 5F *upper panel***). In the placenta, *Cisd1* mRNA expression was increased and *Dpp4* mRNA expression was decreased in DHT+INS-exposed pregnant rats compared to controls (**Fig. 5E *lower panel***). A decrease in placental *Dpp4* mRNA expression was also observed in INS-exposed pregnant rats (**Fig. 5E *lower panel***). However, there was no significant difference in MOTS-c protein levels in the placenta between any of the experimental groups (**Fig. 5F *lower panel***).

### Aberrant regulation of necroptosis-related and anti-/pro-apoptosis-related gene and protein expression

Different types of cell death are seen in uterine and placental tissue during healthy and pathological pregnancy (51-54). To extend our observations on the effect of DHT and INS on ferroptosis and mitochondrial impairment, we analyzed the expression of necroptosis (*Mlkl, Ripk1*, and *Ripk3*), anti-apoptosis (*Bcl2* and *Bcl-xl*) and pro-apoptosis (*Bax, Bak, Casp3*, and cleaved caspase-3) mRNAs and proteins (6, 8, 10) in the gravid uterus and placenta. As shown in **Figure 6A**, DHT+INS-exposure significantly decreased uterine *Ripk1* mRNA expression, while uterine *Mlkl* and *Ripk3* mRNAs were increased by DHT and/or INS exposure when compared to control pregnant rats (**Fig. 6A *upper panel***). Furthermore, co-exposure to DHT and INS increased *Bcl2, Bcl-xl*, and *Bax* mRNA expression in the gravid uterus, with similar increases in these genes seen in DHT-exposed and/or INS-exposed pregnant rats compared to controls (**Fig. 6B *upper panel***). In DHT+INS-exposed pregnant rats, *Casp3* mRNA expression and cleaved caspase-3 protein abundance were decreased in the gravid uterus (**Fig. 6B *upper panel* and C**). In contrast, in the placenta we found that both *Ripk1* and *Ripk3* mRNAs were increased in DHT+INS-exposed pregnant rats compared to controls (**Fig. 6A *lower panel***). Furthermore, maternal co-exposure to DHT and INS increased placental *Bcl2, Bcl-xl, Bax*, and *Bak* mRNA expression (**Fig. 6B *lower panel***). There were, however, no changes in *Casp3* mRNA expression or cleaved caspase-3 protein abundance in the placenta (**Fig. 6B *lower panel* and C**). Lastly, similar increases in placental *Bcl2, Bcl-xl, Bax, Bak* and *Casp3* mRNAs were seen in DHT-exposed and/or INS-exposed pregnant rats compared to controls (**Fig. 6B *lower panel***).

**Figure 6.**
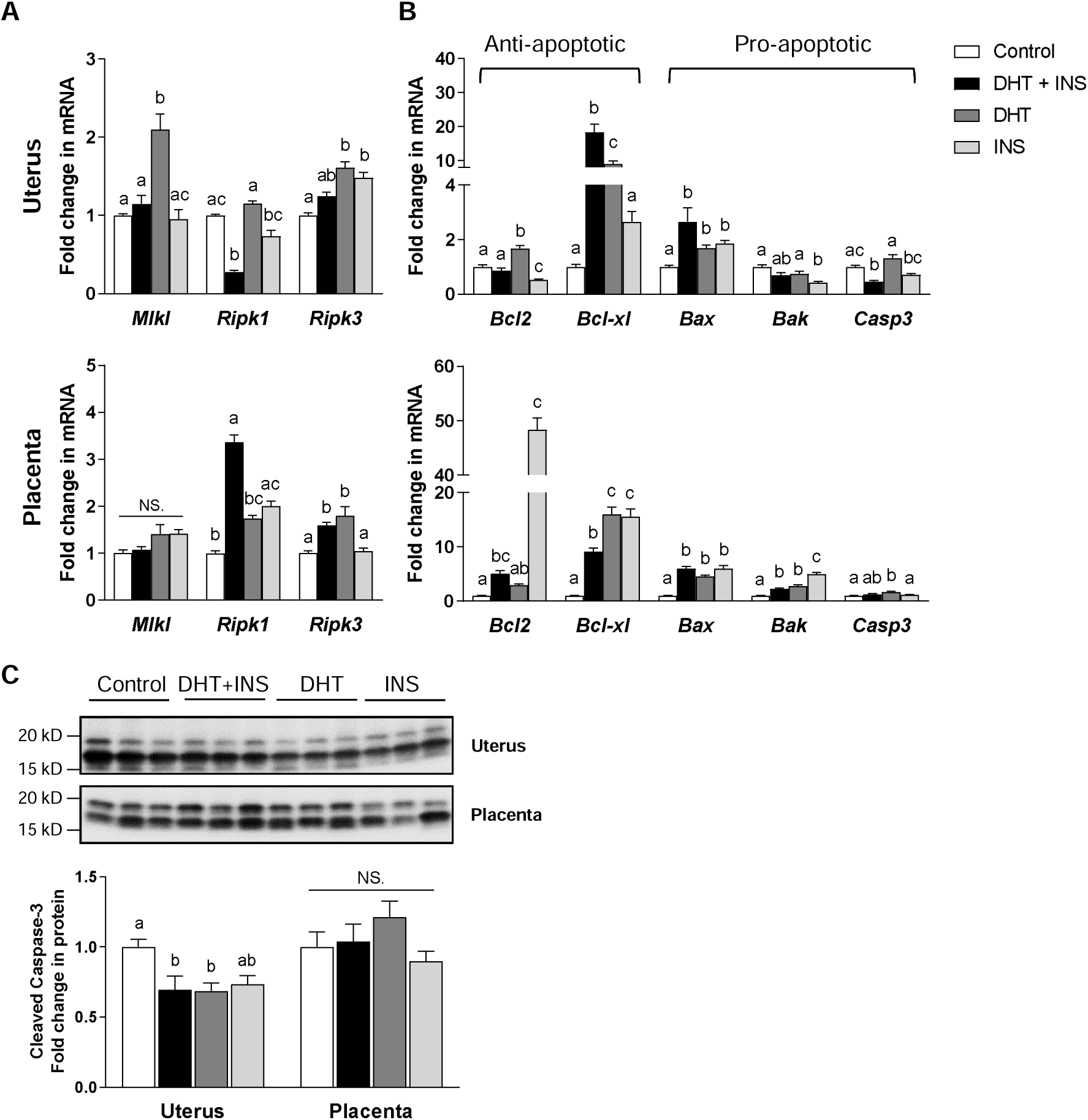
The regulatory pattern of necroptosis-related and pro-/anti-apoptosis-related gene and protein expression in pregnant rats exposed to DHT and/or insulin at GD 14.5. qPCR analysis of *Mlkl, Ripk1, Ripk3, Bcl2, Bcl-xl, Bax, Bak*, and *Casp3* mRNA in the uterus and placenta (A and B, n = 8/group). Western blot analysis of cleaved caspase-3 protein expression in the uterus and placenta (C, n = 9/group). In all plots, values are expressed as means ± SEM. Significant differences (p < 0.05) within each group are denoted by different letters, and the same letter between groups indicates lack of statistical significance. N.S., not significant. DHT, 5α-dihydrotestosterone; INS, insulin.

## Discussion

Because PCOS patients frequently suffer from miscarriage and infertility (3, 4), it is important to understand the molecular mechanisms through which HAIR affects tissues such as the gravid uterus and placenta. Until now, there have been no reports exploring the relationship between PCOS and regulated cell death in the uterus and placenta. Our results thus fill an important clinically relevant knowledge gap by experimentally demonstrating that maternal HAIR can cause the activation of ferroptosis in the gravid uterus and placenta, although this is mediated through different molecular and cellular mechanisms. Alterations in the ferroptosis pathway in the uterus and placenta as a result of maternal HAIR likely contribute to impaired fetal survival seen in experimental animal models, as well as women with PCOS.

Mammalian Gpx4 generally functions as a major antioxidant regulator of systemic and cellular responses to oxidative stress, and loss of function of Gpx4 protein and depletion of GSH levels are the key mechanisms for triggering ferroptosis (7, 11). *In vivo* knockout studies have shown that mice lacking the entire *Gpx4* gene are early embryonic lethal (55), and Gpx4-deficient male mice are completely infertile (56). Although the presence of Gpx4 has been shown in uteri from cows (57, 58) and pigs (59), the physiological role and localization of Gpx4 have not been demonstrated in human and rodent reproductive tissues, including the uterus. Here, we show that Gpx4 is widely expressed in the human and rat uterus, including by decidualized stromal cells. Furthermore, Gpx4 is down-regulated in the gravid uterus by maternal exposure to DHT and INS. Correspondingly, the levels of GSH are decreased and GSH+GSSG levels are increased in the uterus by DHT and INS co-exposure. Taken together, these results suggest that maternal HAIR disrupts the Gpx4-GSH regulatory axis and can result in the induction of ferroptosis in the uterus during pregnancy. The finding that GSH levels in the uterus were lowest in the INS-only treated rats that showed a non-significant reduction in Gpx4 suggests that other pathways and factors such as the antioxidants Nrf1 and Nrf2 (60) might be altered by the treatments and contribute to the resultant changes in GSH status and should be investigated for causality in the future (also in the placenta of DHT and/or INS-exposed dams). Indeed, we have previously found altered abundance of antioxidants in pregnant rats exposed to DHT and/or INS (18, 19). Consistent with previous work on the human placenta (61, 62), the present study shows that the Gpx4 protein is highly expressed in the rat placenta during pregnancy. Although analysis of whole placental homogenates showed no significant decrease in Gpx4 levels, immunolocalization revealed loss of Gpx4 in specific cell types in the placenta (the glycogen and spongiotrophoblast cells) in response to maternal co-exposure to DHT and INS. The more minor alterations in Gpx4 abundance, combined with the high levels of GSH+GSSG and absence of changes in GSH levels in the placenta, suggest that maternal HAIR induces ferroptosis to a lesser extent in the placenta compared to the gravid uterus. Gpx4 is known to protect cells/tissues against lipid peroxidation by inhibiting lipid-associated hydroperoxides (11). In addition, genetically ablating or inducing decreased Gpx4 expression leads to ferroptosis (14, 57). Taken together, our data therefore suggest that HAIR-induced ferroptosis is mediated by both dysregulation of Gpx4 expression and aberrant increases in lipid peroxidation. The induction of uterine and placental ferroptosis by maternal exposure to DHT and INS may be a novel mechanism contributing to the malfunction of those tissues and hence impaired fetal development during pregnancy. However, how maternal HAIR-mediated uterine and placental ferroptosis compromises the growth and development of the fetus is not clear at this time and should be the subject of future investigations. Moreover, future work should be employed to assess whether the activation of ferroptosis, lipid peroxidation and poor fetal outcomes by maternal HAIR may be preventable by antioxidant administration.

Iron can serve as an essential signaling molecule that modulates diverse physiological processes, and iron homeostasis is required for the normal growth and development of the placenta and fetus during pregnancy (13, 63). An extensive body of evidence indicates that while iron deficiency is linked to abnormal pregnancy (13) and increased risk of fetal death (64), iron overload is associated with the manifestation of PCOS (65). Previous findings by Kim and colleagues indicate that increased circulating iron levels are associated with metabolic abnormalities, including HAIR, in PCOS patients (66). Several studies have demonstrated that in addition to its antioxidative property, Ho1 is a critical regulator for mobilization of intracellular pools of free iron (67, 68). More recently, we have demonstrated that maternal co-exposure to DHT and INS suppresses *Ho1* mRNA expression in the gravid uterus, but not in the placenta (18, 19). While the uptake of transferrin-bound iron, a major maternal iron source for placental transfer, is mainly mediated through iron import proteins such as transferrin receptor 1 (Tfr1, Tfrc) (63), our results show that combined exposure to DHT and INS increases *Tfrc* mRNA expression in association with increased iron deposition in the gravid uterus. Further, we have provided ultrastructural evidence that shrunken mitochondria with numerous electron-dense cristae, a key feature of ferroptosis-related mitochondrial morphology are present in the gravid uterus. However, the placentas of the same animals exhibited increased mRNA expression of *Cisd1*, a mitochondrial iron export factor, but no change in *Tfrc* mRNA or iron accumulation. Ferroptosis can be induced by excessive accumulation of free iron in tissues and cells (69) and our findings support the notion that, in response to exposure to DHT and INS, aberrant iron accumulation and activation of ferroptosis occurs in the gravid uterus but not in the placenta. The synthesis of heme and iron-sulfur clusters is controlled by different intracellular compartments, including the cytosol and mitochondria (69). Further investigations are needed to determine which cellular compartments contribute to the defective utilization of iron and increased ferroptosis observed in the gravid uterus under conditions of HAIR.

Given that aberrant accumulation of intracellular iron induces oxidative stress (69) and subsequently results in multiple modes of cell death (70), it is not surprising that, in addition to ferroptosis, apoptosis (a non-inflammatory form of cell death) and necroptosis (a pro-inflammatory form of cell death) may also be involved in HAIR-induced fetal loss in pregnant rats. Indeed, pregnant rats co-exposed to DHT and INS exhibited decreased *Casp3* mRNA expression and cleaved caspase-3 protein abundance in the uterus, but not in the placenta, even though selectively increased expression of anti-apoptotic genes (*Bcl2* and *Bcl-xl*) and pro-apoptotic genes (*Bax*) was observed in both tissues. We suspect that suppression of apoptosis might serve as a compensatory mechanism to protect against increased ferroptosis in order to maintain homeostasis of the gravid uterus after exposure to DHT and INS. This is supported by studies assessing the interaction and interplay of different cell death pathways in cancer research (71). Ferroptosis and necroptosis are two different forms of regulated necrosis (8, 10). Necroptosis requires mitochondrial ROS generation and is primarily regulated by the Ripk1, Ripk3, and Mlkl proteins (9, 46). We found that in DHT+INS-exposed pregnant rats the level of ROS (19) and expression of *Ripk1* and *Ripk3* mRNAs was increased in the placenta, but not in the gravid uterus ((18) and this study). Therefore, it is tempting to speculate that the activation of necroptosis in response to PCOS-related HAIR might serve to counteract the ferroptosis pathway in the placenta. Additionally, both ferroptosis and necroptosis might intersect and crosstalk with HAIR-induced oxidative damage and subsequently result in increased fetal loss. It remains to be determined whether HAIR-induced pregnancy loss is due to increased iron-mediated uterine ferroptosis or to necroptosis-related defects in the placenta, or both.

In this study, we found that some pro-ferroptosis genes such as *Acsl4, Tfrc* and *Dpp4* were oppositely regulated in the uterus by co-exposure to DHT and INS. However, several anti-ferroptosis genes, including *Slc7a11, Gcls*, and *Cisd1* were downregulated in the gravid uterus after co-exposure to DHT and INS. These results suggest that the suppression of anti-ferroptosis gene transcription might play a dominant role in promoting ferroptosis in this tissue under conditions of HAIR. Compared to the gravid uterus, the placenta showed a distinct profile of ferroptosis-related gene changes in response to the combined DHT and INS exposure. Furthermore, we often observed contrasting expression patterns of pro- and anti-ferroptosis genes in the gravid uterus and placenta with exposure to DHT or INS alone compared to the combined exposure. We do not know the exact reason for these inconsistencies; however, we do know that the expression of ferroptosis-related genes and proteins are only assessed at one gestational age in pregnant rats when they display HAIR (18, 19). In addition, perhaps components of HAIR may act synergistically or through separate pathways to bring about divergent effects on gene expression and signaling pathways to regulate the ferroptosis process. Overall, our findings demonstrate the complexity and challenges in establishing direct roles and patterns linking individual pro-/anti-ferroptosis genes to the ferroptosis pathway in the gravid uterus and placenta in response to DHT and/or INS *in vivo*. Future work should therefore investigate the tissue-specific and time-dependent changes in ferroptosis-related gene expression in the gravid uterus and placenta during the hormonal manipulation. In comparison to the single treatment groups (DHT or INS), specific changes within the maternal uterus and placenta appeared to be driven by hyperandrogenism, insulin resistance, or both (co-treatment with DHT and INS) and reflected the complexity in working with tissues from animals in which many physiological parameters may be altered. Indeed, experiments utilizing gene and pathway inhibitors in uterine decidual cells and placental trophoblasts would be beneficial in future studies that aim to explore the causality of changes observed regarding ferroptosis and iron metabolism. Work is also required to assess whether elevated ferroptosis in the uterus contributes to the placental dysfunction in the rat dams with HAIR due to DHT and INS, which would be aided by a time-course analysis.

Recently, Zhang and colleagues reported that oxidative stress-induced ferroptosis contributes to the pathogenesis of preeclampsia (72). Because decreased Gpx4, GSH, and SLC7A11 protein levels and increased MDA content are seen in the preeclamptic placenta in humans and rats (72), our findings together with this report support the notion that defective ferroptosis is involved in the pathophysiological processes of female reproductive disorders.

In summary, our findings suggest maternal exposure to DHT and INS alters the ferroptosis pathway in the gravid uterus and placenta; however, this occurs via different regulatory mechanisms and signaling pathways. For instance, in contrast to the placenta, increased ferroptosis in the gravid uterus in response to DHT and INS was related to decreased Gpx4 and GSH abundance, altered expression of ferroptosis-associated genes (*Acsl4, Tfrc, Slc7a11*, and *Gclc*), increased MDA and iron deposition, upregulation of the ERK/p38/JNK pathway and mitochondrial *Dpp4* expression, and the appearance of typical ferroptosis-related mitochondrial morphology. In addition, DHT and INS were associated with reduced activation of apoptosis in the uterus and increased necroptosis in the placenta. The concomitant presence of different forms of regulated cell death would be expected to disrupt uterine and placental function and play a role in the fetal loss observed in DHT+INS-exposed pregnant rats. Both the maternal uterine decidua and placenta play essential roles in embryo implantation and successful pregnancy (73, 74). Therefore, while the present study improves our understanding of the impact of HAIR on regulated cell death in specific tissues during pregnancy, more preclinical and clinical studies are needed to further investigate the molecular and functional connectivity between the maternal decidua and the placenta and between the placenta and fetus under conditions of PCOS.

## Abbreviations

PCOS: polycystic ovary syndrome;
HAIR: hyperandrogenism and insulin resistance;
DHT: 5α-dihydrotestosterone;
INS: human recombinant insulin;
Gpx4: glutathione peroxidase 4;
GSH: glutathione (reduced state);
GSSG: glutathione disulfide (oxidized state);
JNK: c-JUN NH2-terminal kinase;
MAPK: mitogen-activated protein kinase;
MDA: malondialdehyde;
GD: gestational day;
TEM: transmission electron microscopy.

## Author contributions

Study design and supervision: LRS. Study conduct: YZ, MH, WJ, GL, JZ, BW, PC, XL, YH, LS, XW, and LRS. Data collection: YZ, MH, WJ, GL, JZ, BW, JL, XL, and LRS. Data analysis: YZ, MH, JL, and LRS. Data interpretation: SL, ANS, LS, MB, LRS, and HB. Drafting the manuscript: YZ, MH, and LRS. Revising the manuscript: SL, ANS, MB, LRS, and HB. YZ, MH, LRS, and HB take responsibility for the integrity of the data analysis. All authors have read and approved the final version of the manuscript.

## Acknowledgments

This study was financed by grants from the Swedish Medical Research Council (grant number 10380), the Swedish state under the agreement between the Swedish government and the county councils – the ALF-agreement (grant number ALFGBG-147791), Jane and Dan Olsson’s Foundation, the Knut and Alice Wallenberg Foundation, and the Adlerbert Research Foundation to HB and LRS as well as the National Natural Science Foundation of China (Grant No. 81774136), the Project of Young Innovation Talents in Heilongjiang Provincial University (Grant No.UNPYSCT-2015121), the Scientific Research Foundation for Postdoctoral Researchers of Heilong Jiang Province, the Project of Science Foundation by Heilongjiang University of Chinese Medicine, and the Project of Excellent Innovation Talents by Heilongjiang University of Chinese Medicine to YZ. The Guangzhou Medical University High-level University Construction Talents Fund (grant number B185006010046) supported MH. ANSP is supported by a Royal Society Dorothy Hodgkin Research Fellowship. The funders had no role in the design, data collection, analysis, decision to publish, or preparation of the manuscript.

