## Supplementary material for "Induction of hyperandrogenism and insulin resistance differentially modulates ferroptosis in uterine and placental tissues of pregnant rats": Suppl Figures 1-5

### Supplemental Figure 1

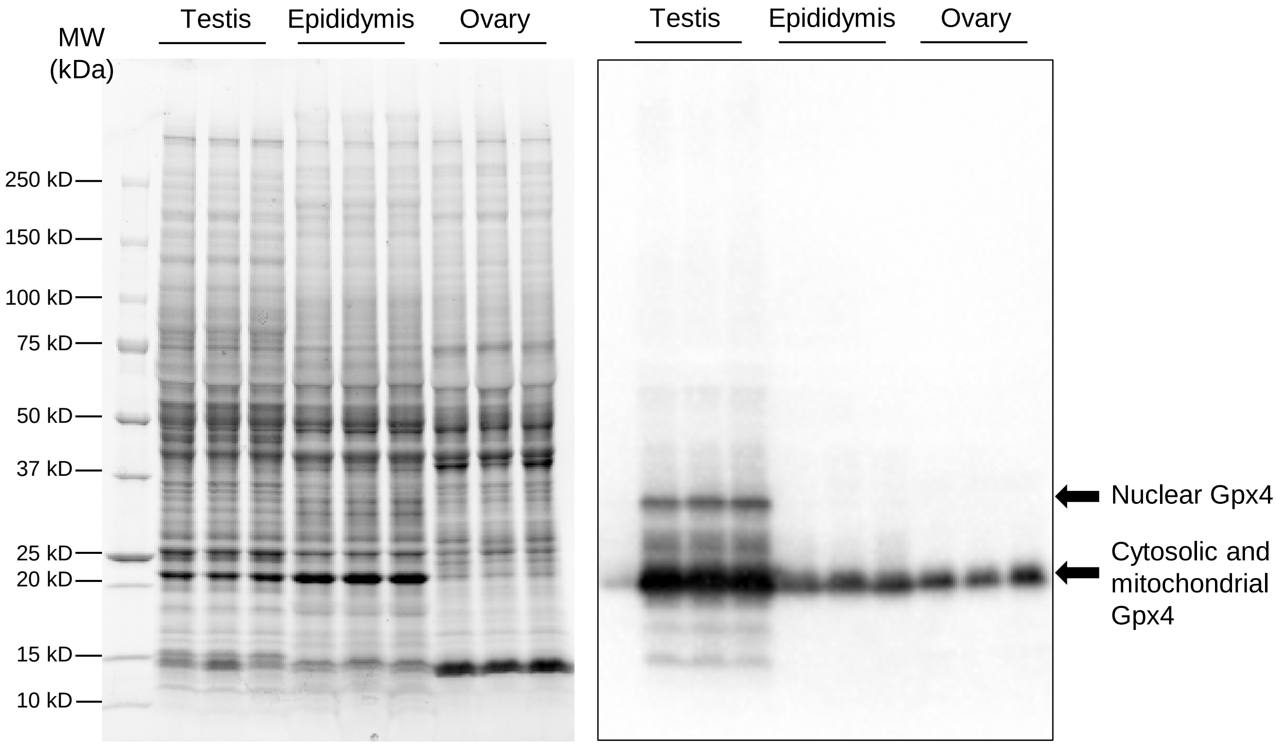


**Supplemental Figure 1. Tissue expression of Gpx4 protein in the adult rats.** Protein samples (*left*) were isolated from selective tissues (testis, epididymis, and ovary; n = 6/tissue). The level of Gpx4 protein expression was determined by Western blot analysis (*right*). Each lane corresponds to the different rat samples. Relative mobilities of molecular mass standards (MW) is shown (in kilodaltons) on the left.

#### Supplemental Figure 2

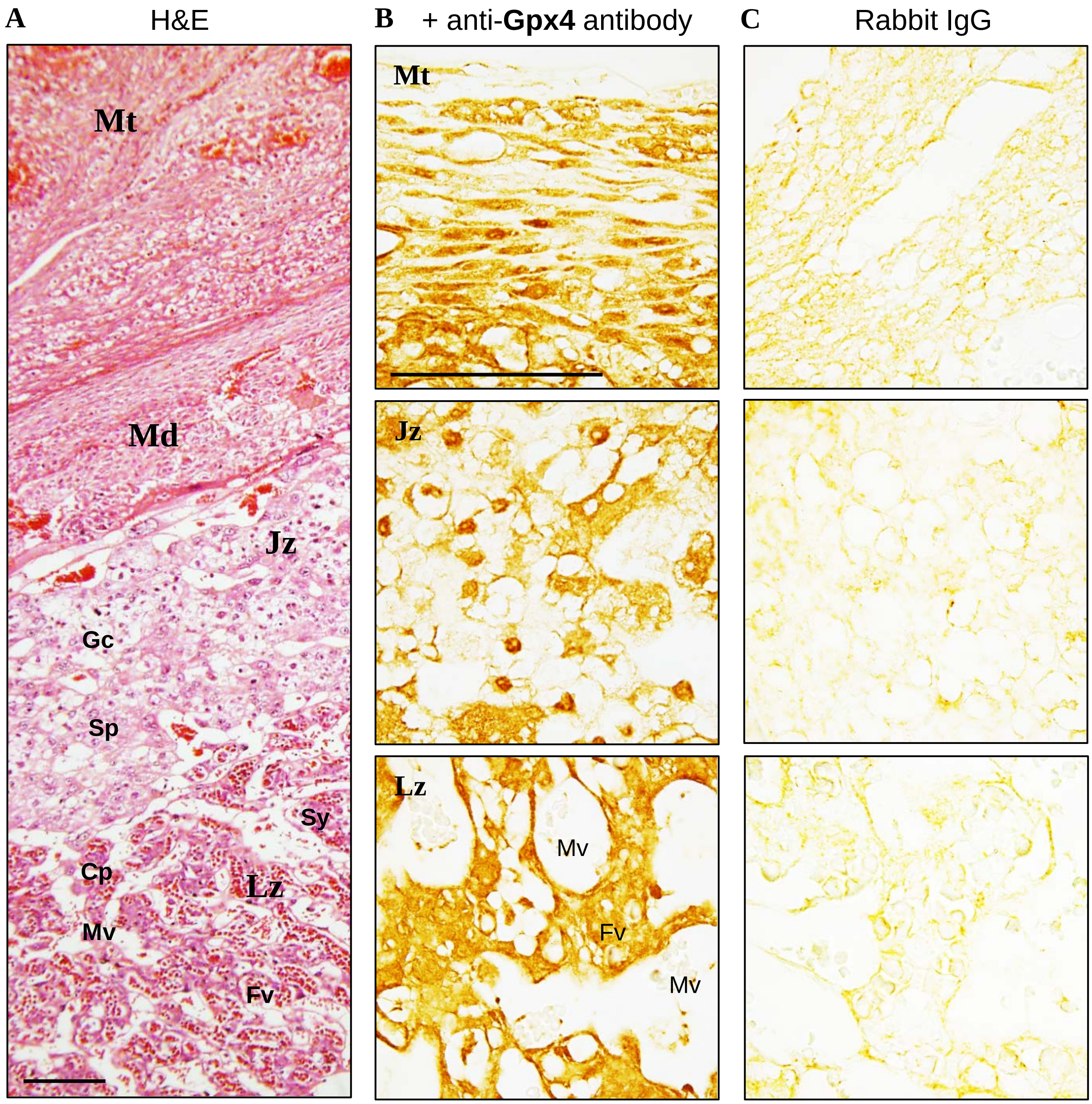


**Supplemental Figure 2. Demonstration of Gpx4 antibody specificity.** Tissue sections were stained with hematoxylin and eosin (A). An antibody against Gpx4 (ab125066, 1:200, Abcam) that reacts with both rat and human tissues was used to determine the immuno-localization of Gpx4 in the rat placenta at GD 14.5. The rat placenta is compsoed of the maternal (Mt and Md) and fetal (Jz, Lz and yolk sac) parts. (B). The same concentration of rabbit IgG instead of the primary and secondary antibodies was used as the negative control (C). Images are representative of 4–6 tissue replicates per group. Mt, mesometrial triangle; Md, mesometrial decidua; Jz, junctional zone (maternal side); Lz, labyrinth zone (fetal side); Gc, glycogen cells; Sp, spongiotrophoblast cells; Cp, cytotrophoblast cells; Sy, syncytiotrophoblast cells; Mv, maternal blood vessel; Fv, fetal blood vessel. Scale bars (100 μm) are indicated in the photomicrographs.

### Supplemental Figure 3

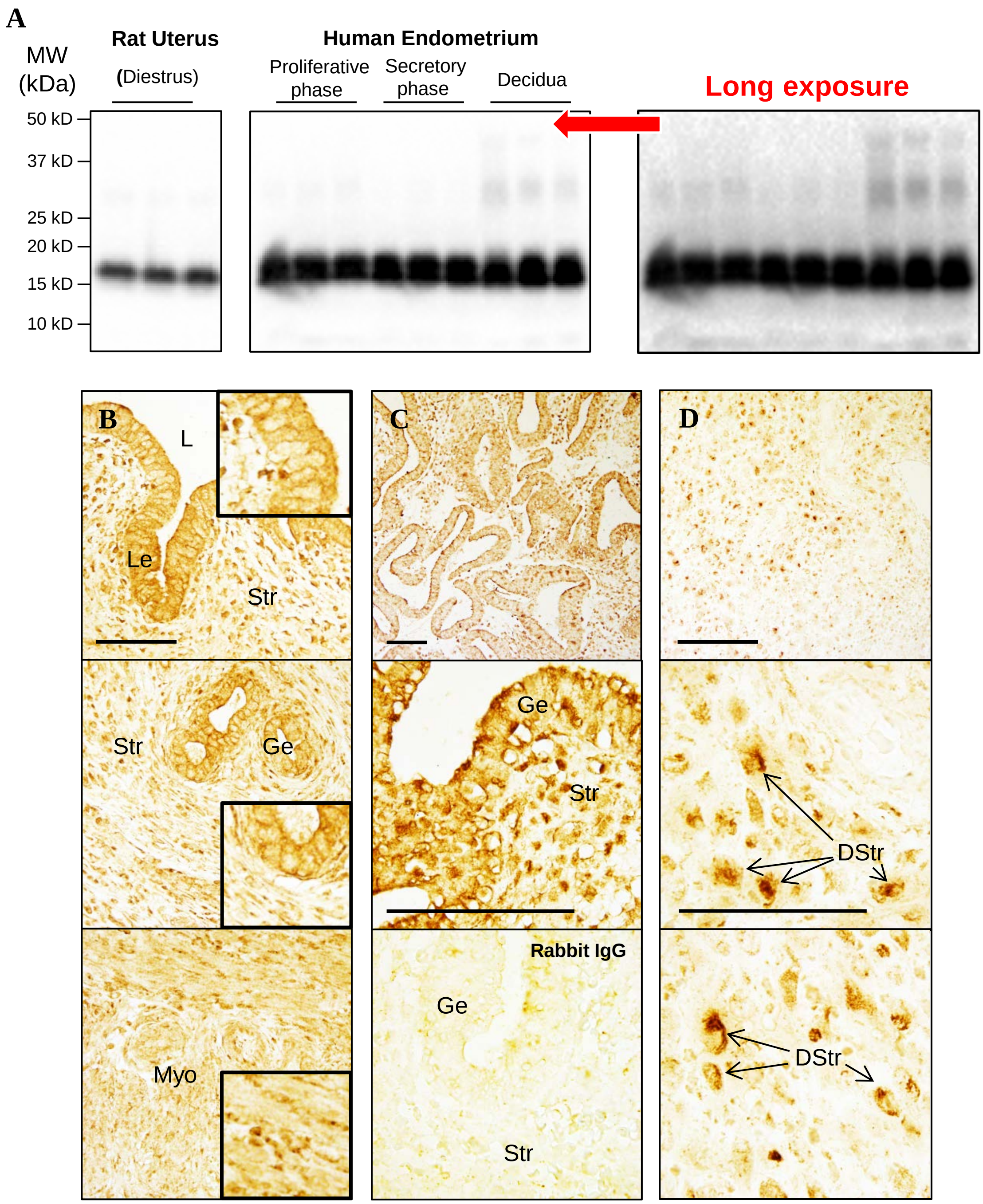


**Supplemental Figure 3. Tissue expression of Gpx4 protein in the adult rat uterus and human endometrium.** The level and localization of uterine Gpx4 protein expression in the diestrus stage in rats and in the non-pregnant proliferative/secretory phase and early pregnant decidualized (the first trimester) endometria in humans were determined by Western blot (A) and immunohistochemical analyses (B–D). In the Western blot analysis, each lane corresponds to the different rat and human samples. Relative mobilities of molecular mass standards (MW) are shown (in kilodaltons) on the left. Immunohistochemical images are representative of 3−5 tissue replicates per group. Detailed views of the boxed areas are shown in the inset (B). Arrows point to the endometrial stromal cells. The same concentration of rabbit IgG instead of the primary and secondary antibodies was used as the negative control. L, lumen; Le, luminal epithelial cells; Ge, glandular epithelial cells; DStr, decidualized stromal cells; Str, stromal cells; Myo, myometrium. Scale bars (100 μm) are indicated in the photomicrographs.

### Supplemental Figure 4

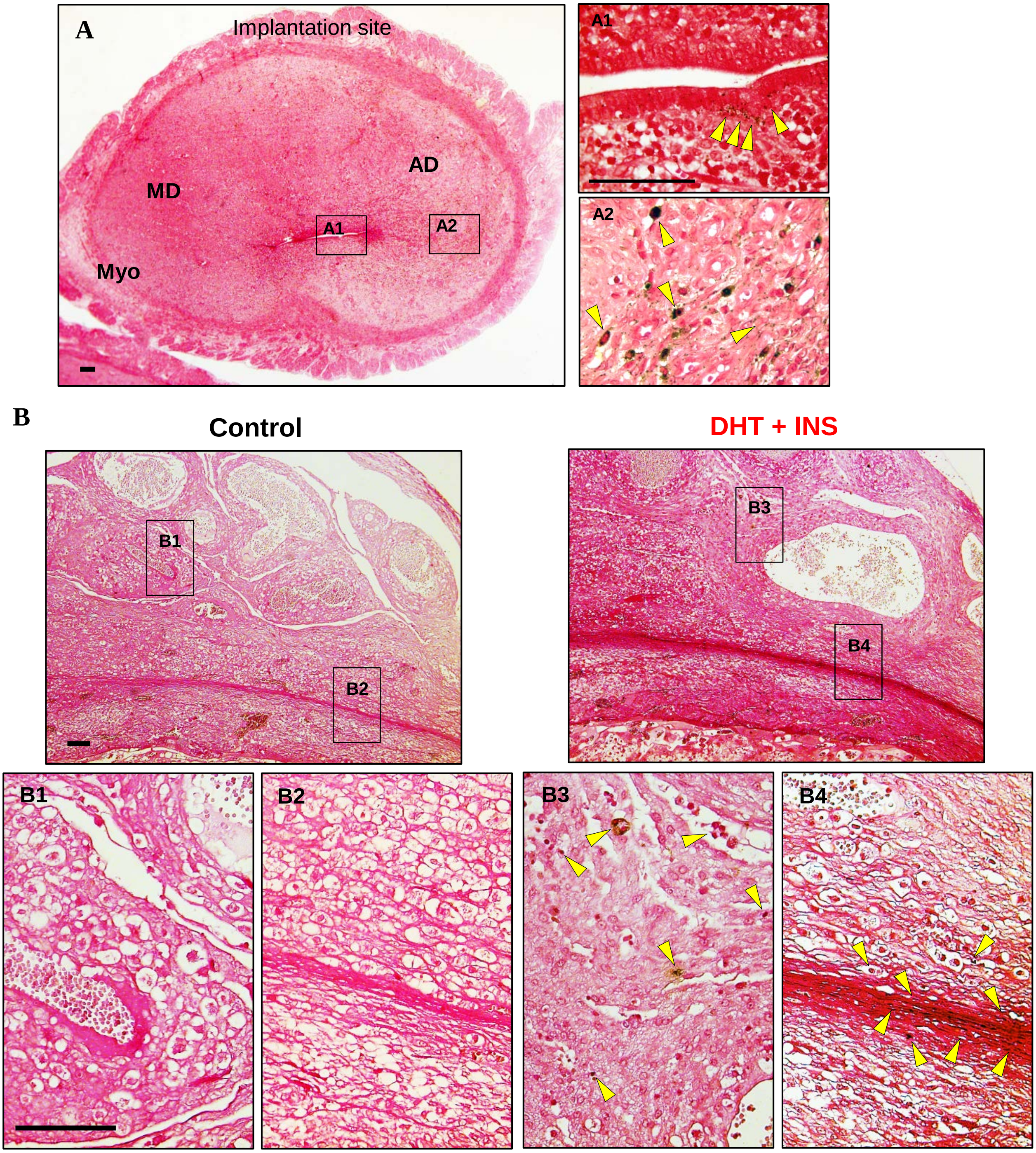


**Supplemental Figure 4. Iron deposition in the uterine sections from pregnant rats at GD 6 (A, A1-2) and pregnant rats treated with vehicle and DHT+INS at GD 14.5 (B, B1-4).** The sections were stained by DAB-enhanced Perls' staining for iron accumulation. Images are representative of 5−8 tissue replicates per group. Yellow arrowheads indicate punctate cytoplasmic, and granular iron-positive staining. MD, mesometrial decidua; AD, antimesometrial decidua; Myo, myometrium. Scale bars (100 μm) are indicated in the photomicrographs. DHT, 5α-dihydrotestosterone; INS, insulin.

#### Supplemental Figure 5

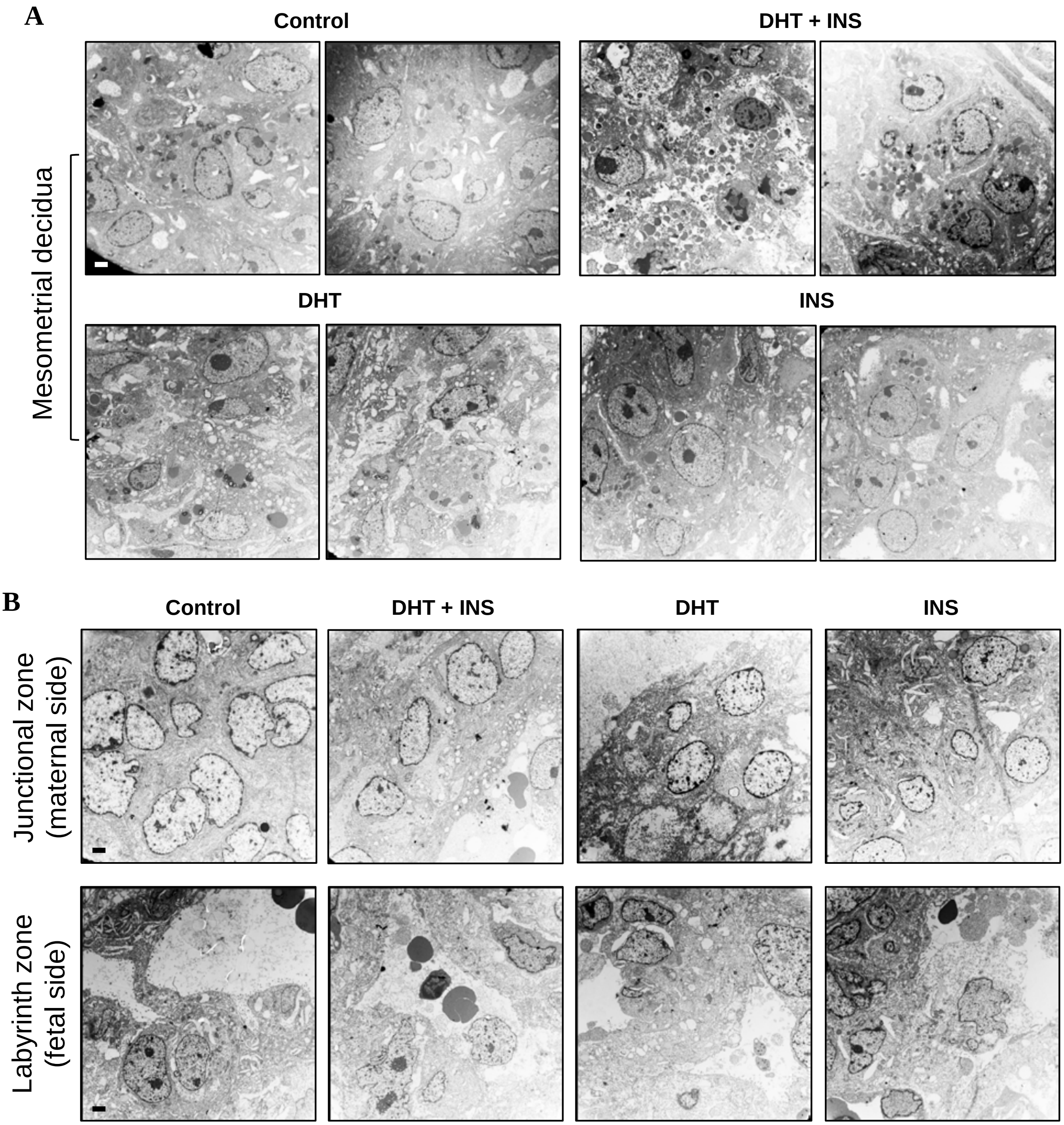


**Supplemental Figure 5. Mitochondrial ultrastructural defects in mesometrial decidua, and basal and labyrinth zones in pregnant rats treated with DHT and/or insulin at GD 14.5.** Scale bars (2 μm) are indicated in the photomicrographs. DHT, 5α-dihydrotestosterone; INS, insulin.

### Supplemental Table 1

| | The statistic model | | | |
|---|---|---|---|---|
| | One way ANOVA followed by Turkey Post-hoc test | | Nonparametric tests followed by Kruskal-Wallis test | |
| **Tissue** | **Uterus** | **Placenta** | **Uterus** | **Placenta** |
| **Protein (n = 9/group)** | GSH, GSH+GSSG (Fig. 2A) | GSH, GSH+GSSG (Fig. 2A) | Gpx4 (Fig. 1A) | Gpx4 (Fig. 1A) |
| | MDA (Fig. 2C) | | | MDA (Fig. 2C) |
| | ERK1/2, p-ERK1/2 and the ratio (Fig. 4A) | ERK1/2, p-ERK1/2 and the ratio (Fig. 4B) | p38, p-p38, and the ratio (Fig. 4A) | p38, p-p38, and the ratio (Fig. 4B) |
| | | | JNK, p-JNK and the ratio (Fig. 4A) | JNK, p-JNK and the ratio (Fig. 4B) |
| | | | MOTS-c (Fig. 5F) | MOTS-c (Fig. 5F) |
| | | | Cleaved Caspase-3 (Fig. 6C) | Cleaved Caspase-3 (Fig. 6C) |
| **Gene (n = 7–8/group)** | Slc7a11 (Fig. 2B) | Slc7a11 (Fig. 2B) | Slc1a5, Acsl4, Gls2, Cs, Gclc, Gss, Tfrc, Ireb2 (Fig. 2B) | Slc1a5, Acsl4, Gls2, Cs, Gclc, Gss, Tfrc, Ireb2 (Fig. 2B) |
| | Cisd1 (Fig. 5E) | | Dpp4 (Fig. 5E) | Cisd1, Dpp4 (Fig. 5E) |
| | Mlkl (Fig. 6A) | Mlkl (Fig. 6A) | Ripk1, Ripk3 (Fig. 6A) | Ripk1, Ripk3 (Fig. 6A) |
| | Bcl2, Bcl-xl, Bak (Fig. 6B) | Bcl-xl, Bax (Fig. 6B) | Bax, Casp3 (Fig. 6B) | Bcl2, Casp3 (Fig. 6B) |
